# Mechanistic insights into the activity of Benzoxaboroles against *Cryptosporidium parvum*: CPSF3 targeting and resistance implications

**DOI:** 10.1101/2025.10.09.681364

**Authors:** Janine Wenker, Laurence Braun, Christopher Swale, Axel Gouin, Julien Pichon, Laura Sedano, Thomas Baty, Irina Dobrescu, Marco Lalle, Mohamed-Ali Hakimi, Alexandre Bougdour, Fabrice Laurent

## Abstract

Benzoxaboroles are emerging as promising treatments against a broad range of protozoan parasites, including *Trypanosoma*, *Leishmania*, *Plasmodium*, *Toxoplasma*, and *Cryptosporidium*. We previously demonstrated that the benzoxaborole compound AN3661 inhibits the endonuclease activity of CPSF3 in *Cryptosporidium hominis*, likely by disrupting pre-mRNA processing, which in turn limits parasite growth. In this study, we further explored its mode of action and found that a specific mutation (Y385N) in CPSF3 of *C. parvum* confers strong resistance to AN3661. Interestingly, this mutation does not affect the parasite’s sensitivity to two other benzoxaboroles, AN13762 and AN7973, which are also thought to target *Cp*CPSF3. All three compounds interfered with mRNA processing in *C. parvum*, consistent with inhibition of CPSF3 complex activity, and showed parasiticidal activity, especially during the late stages of merogony, blocking parasite egress. Two of them also impaired gamogony. In *T. gondii*, AN7973 remained effective against several strains carrying mutations within TgCPSF3 that confer resistance to AN3661 and AN13762, suggesting that it might represent an alternative chemotype targeting CPSF3 with the potential to overcome resistance. Notably, AN7973 successfully controlled severe infection in susceptible mice challenged with the AN3661-resistant *C. parvum* strain carrying the CpCPSF3_Y385N_ mutation. Finally, we extended the known antiparasitic spectrum of AN3661 and AN7973 to include *Eimeria tenella* and *Giardia duodenalis*, two important pathogens in veterinary and human health. Altogether, our findings refine the understanding of CPSF3-targeting benzoxaboroles, identify alternative chemotypes with the potential to bypass resistance, and support their potential use in combination therapies to delay or prevent the emergence of drug resistance.

**Author summary:** Protozoa are single-celled parasites that cause significant morbidity and mortality worldwide, affecting both humans and animals. The development of new targeted therapies with highly selective compounds requires a detailed understanding of their mode of action and precise interactions with parasite targets. Benzoxaboroles are a potent class of molecules active against *Cryptosporidium*, with compound AN3661 known to inhibit the endonuclease activity of the Cleavage and Polyadenylation Specificity Factor 3 (CPSF3) in *Cryptosporidium hominis*. Here, we show that this inhibitory effect is critical during both asexual and sexual developmental stages and that tyrosine at position 385 of *C. parvum* CPSF3 plays a key role in its inhibition. A mutation at this site confers resistance to AN3661 both *in vitro* and *in vivo*, but not to two other benzoxaboroles, AN13762 and AN7973. Notably, all three compounds disrupted pre-mRNA processing in *Cryptosporidium*, consistent with inhibition of the CPSF3 complex. Using *Toxoplasma*, a related protozoan that allows more efficient genetic manipulation, we found that none of the six mutations conferring resistance to compounds AN3661 and AN13762 conferred resistance to compound AN7973. AN7973, which strongly inhibits both *C. parvum* and *T. gondii*, may therefore represent an alternative chemotype targeting CPSF3. Furthermore, we demonstrated that the antiparasitic spectrum of AN3661and AN7973 extends to include *E. tenella* and *G. duodenalis*, two important pathogens in veterinary and human health. Altogether, our results refine the understanding of three CPSF3-targeting benzoxaboroles in *Cryptosporidium* and identify compound AN7973 as capable of overcoming resistance to AN3661, thereby supporting the rationale for combination therapies to prevent the emergence of drug resistance.

## Introduction

Protozoan diseases cause significant health and economic burdens worldwide, particularly in low-income regions. In humans, they are responsible for high morbidity and mortality, while in animals they impair health and reduce productivity. Many of these diseases are zoonotic, impacting both public health and animal species. Their control is challenged by emerging drug resistance, often by the absence of effective vaccines, and the limited availability of new therapies. It is therefore essential to deepen our understanding of these diseases, investigate resistance mechanisms, develop new treatments, and adapt control strategies.

*Cryptosporidium* exemplifies these challenges. Highly infectious and resistant to chlorine, it can trigger large waterborne outbreaks. In humans, it causes severe, persistent diarrhea in young children and immunocompromised individuals, leading to malnutrition, developmental delays, and substantial mortality in low-resource settings [1]. In livestock, it is a leading cause of neonatal diarrhea in ruminants, driving dehydration, poor growth, mortality, and major economic losses [2, 3]. Infected animals also act as reservoirs, fueling zoonotic transmission and complicating control efforts. Improved diagnostics and integrated control strategies that consider human, animal, and environmental health are essential. New drugs are urgently needed, as current treatments are only partially effective, especially for vulnerable populations such as young children, immunocompromised patients (e.g., HIV/AIDS), and malnourished individuals [4, 5]. In livestock, available treatments require preventive administration and do not fully clear infections. Developing safe and effective therapies for both humans and animals is crucial to enhance health and welfare, reduce economic impact, and break the cycle of transmission between species.

Benzoxaboroles have emerged as promising compounds against several protozoan parasites, including *Trypanosoma*, *Leishmania*, *Plasmodium*, *Toxoplasma*, and *Cryptosporidium* [6–15]. Notably, we previously demonstrated that AN3661 can inhibit the endonuclease activity of *C. hominis* CPSF3 and proposed that this effect may underlie interference with its pre-mRNA processing function, potentially accounting for the observed reduction in parasite growth following treatment [6]. Here, we aimed to build on this initial model by confirming that one of the main CPSF3 amino acid variants— Y408S in *Plasmodium* and Y483N in *Toxoplasma*—can also confer resistance to AN3661 when introduced into *C. parvum* CPSF3. We also investigated the role of two other benzoxaboroles, AN13762 and AN7973, which have been shown to block *C. parvum* development and are suggested to interact with CPSF3 [10, 14]. We recently showed that AN13762 also inhibits CPSF3 in *T. gondii* and represents an alternative chemotype to AN3661; however, its mechanism of action against *Cryptosporidium* has yet to be elucidated. AN7973 is a potent inhibitor of *Cryptosporidium in vitro* and *in vivo*. All these compounds possess a conserved boron-containing heterocycle that is critical for their biological activity.

In this study, we found that benzoxaborole compounds exert parasiticidal activity by targeting the late stages of *Cryptosporidium* merogony and gamogony. The key mutation Y385N in CPSF3 conferred strong resistance to AN3661, but not to AN7973 or AN13762. In *T. gondii*, AN7973 is active, and none of the mutations that conferred resistance to AN3661 and/or AN13762 provided increased resistance to AN7973. However, importantly, all three compounds disrupted mRNA processing in *C. parvum*, suggesting that AN13762 and AN7973 act through the same molecular target, CPSF3, while representing alternative chemotypes to AN3661. This distinction is particularly relevant for limiting the emergence of resistance mechanisms *in vivo*. For instance, AN7973 effectively controlled infection in susceptible mice, heavily infected with an AN3661-resistant (Y385N) *C. parvum* strain. Furthermore, we expanded the spectrum of protozoan parasites susceptible to AN3661 and AN7973 to include *Eimeria tenella* and *Giardia duodenalis*, pathogens of veterinary and human health concern, respectively. Altogether, our results refine the understanding of the mechanisms of action of AN3661, AN7973, and AN13762, broaden their potential applications, and offer valuable insights for the development of combination therapies aimed at limiting future resistance.

## Results

### AN3661, AN7973 and AN13562 block intracellular schizogony and gamogony but do not interfere with invasion process

We further investigated the inhibitory mechanism of the benzoxaborole compounds AN3661, AN7973 and AN13562, on parasite invasion and intracellular development using a transgenic strain expressing Nluc and mCherry (*Cp*-INRAE-Nluc-mCh) [16]. Preincubation of HCT-8 cells with the three benzoxaborole compounds before invasion for 3 h prior to invasion did not reduce the invasion process (**Fig 1A**). A comparable lack of effect was also observed when the pre-incubation period was extended to up to 15 h (data not shown). Likewise, their presence during excystation and invasion had no effect on parasite entry or early growth. In contrast, exposure during intracellular development strongly reduced parasite proliferation. Time-course assays revealed that treatment during first-generation schizont maturation was most effective. The 8-12h window was the most critical (**Fig1-B**), coinciding with the intense transcription required to generate the first-generation merozoites.

**Figure 1:**
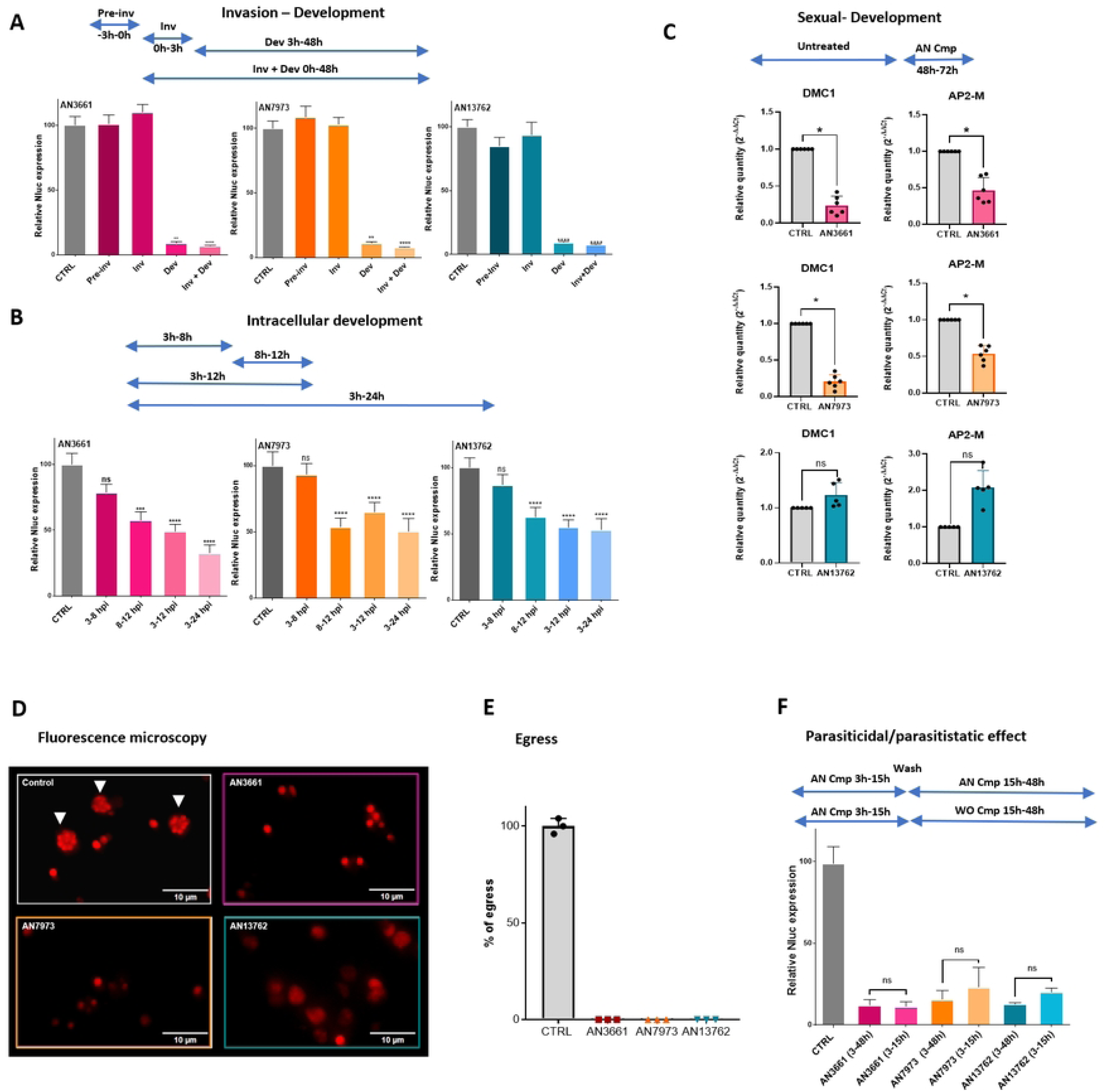
Effect of AN3661, AN7973, and AN13762 on *in vitro* parasite development. Drug were added at different time in cell culture, previous to infection with *Cp*-INRAE-Nluc-mCh strain (A), during the invasion step (A) or during different periods of time during intracellular development (A-B). Parasite development was measured at 48hpi. For drug effect on sexual development drug were added at 40hpi and RNA extracted at 72hpi for Q-RT-PCR analysis (C). Time-lapse microscopy was performed in controlled growth conditions from 5hpi. Single images show parasite development at 12hpi visible in red. White arrows indicate mature meronts visible in control well (D). Video generated with Fiji software were used to evaluate the rate of egress of first generation merozoites (E). The ability of parasites to resume growth after drug removal was assessed by comparing Nluc activity at 48hpi between samples continuously exposed to the drugs and those from which the drugs were removed at 12h post-infection (F). Statistics: Kruskal-Wallis test followed by 1-way Anova multiple comparison **** p<0.0001; *** p<0.001 ** p<0.005 (A, B, F). Wilcoxon matched-pairs signed rank test on n=5-6 independent experiments. * p=0.03 (C).

To evaluate the effect of benzoxaborole compounds on gamogony, we designed primers for RT-qPCR analysis targeting male- and female-specific transcripts. A recent study identified AP2-M gene (cgd6_2670) as being expressed in the sexual development. Using a fluorescent reporter under the control of the AP2-M promoter, Tandel *et al.* [17] demonstrated that AP2-M serves as a male-specific marker. Similarly, DMC1 (cgd7_1690), a homolog of the RecA/RAD51 recombinase family, plays a critical role in meiotic homologous recombination, with a peak expression occurring during the sexual development stages in macrogamonts [18]. When AN3661 and AN7973 were added to the culture medium from 40 hours post-infection (hpi) onward, a marked reduction in the expression of both male and female markers was observed *in vitro (***Fig1-C**). This extended our previous findings on AN3661 and confirmed the initial observations reported by Lunde *et al.* regarding the potency of AN7973 against asexual-to-sexual conversion [14]. In contrast, AN13762, which requires substantially higher concentrations *in vitro* to control asexual development, did not significantly affect sexual development under the tested conditions (**Fig1-C**).

*Cp*-INRAE-Nluc-mCh [16] was used to perform time-lapse microscopy and monitor the egress of first-generation merozoites and subsequent reinvasion events. In HCT-8 cells treated with AN3661, AN7973, or AN13762 at concentrations corresponding to 10 times their respective EC₅₀ values, parasites remained arrested at the trophozoite stage, failing to fully mature compared to untreated controls (**Fig1-D**), and no egress of first generation merozoites was observed (**Fig1-E**). Finally, we assessed whether the drugs exerted parasiticidal or parasitistatic effects by removing the compounds from the culture medium at 16 hours post-infection and monitoring parasite regrowth through Nluc activity at 48 hours post-infection (**Fig1-F**). No recovery occurred, demonstrating parasiticidal activity. These observations align with CPSF3 targeting proposed in earlier studies, prompting us to further investigate the drug-CPSF3 interactions.

### Y385 is a key Amino-Acid for AN3661 but not AN13762 nor AN7973 interactions with *Cp*CPSF3 *in vitro* and *in vivo*

The crystal structure of *Ch*CPSF3 bound to AN3661 revealed several amino acids involved in the interaction mechanism [6]. Among them, key hydrophobic contacts were identified between carbons 3-4 of AN3661 and the aromatic Y385; between carbon-1 and V35; and between the ethyl of AN3661 and F264. In *T. gondii*, we identified single mutants in CPSF3 that were able to induce a strong resistance to AN3661 (Y483N, E545K, Y328C/H) [7] and to AN13762 (G456S, Y483N, E545K, Y328C, Y328H, S519C) [10]. Using a forward genetic approach we demonstrated in *T. gondii that* Y483N was one of the mutations conferring the strongest resistance to both AN3661 and AN13762. Consequently, a transgenic *C. parvum* strain harboring a mutation at residue Y385 was engineered to assess its capacity to confer resistance to the three benzoxaboroles under both *in vitro* and *in vivo* conditions (**Fig2-A**).

**Figure 2:**
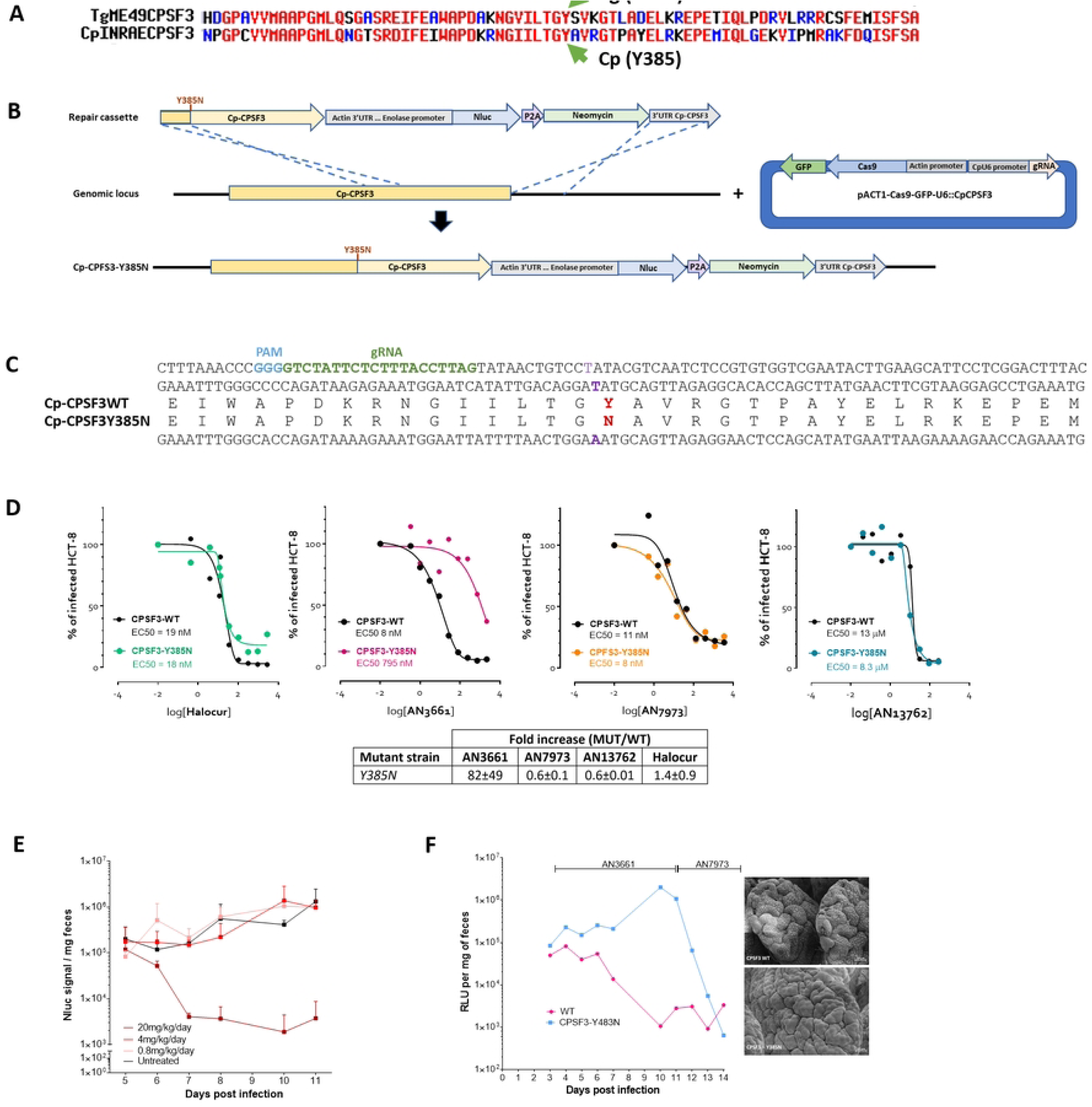
Generation and characterization of the CpY385N strain with *in vitro* and *in vivo* resistance profiling. Partial alignment of CpCPSF3 and TgCPSF3 revealing amino-acid conservation (A). Diagram illustrating the genome-editing strategy used for generating transgenic *C. parvum* line (*Cp*-INRAE-Nluc-CPSF3^Y385N^) expressing the N385 at the endogenous locus (B). Nucleotide sequence of *Cp*-INRAE-Nluc-CPSF3^Y385N^ strain was validated by sequencing. The graphic representation shows the PAM site (highlighted in blue), the gRNA target sequence (highlighted in green), nucleotide substitution (highlighted in purple) conferring the 383N mutation (highlighted in red) (C). EC_50_ determination *in vitro* comparing growth inhibition of *Cp*-INRAE-Nluc-mCh to *Cp*-INRAE-Nluc-CPSF3^Y385N^ in presence of various concentrations of Halocur™, AN3661, AN7973, or AN13762. Table represents fold increase in resistance (± SD) for mutants compared to CPSF3 wild-type strain, in 3 independent experiments (D). Determination of the optimal AN3661 concentration to be administered in the drinking water of GKO mice to control *C. parvum* replication. Four groups of three mice were infected with 5 × 10⁶ *C. parvum* oocysts. Starting on day 3 post-infection, animals from three of the groups received drinking water supplemented with AN3661 at doses of 0.8 mg/kg, 4 mg/kg, or 20 mg/kg, respectively. Parasite excretion was monitored daily by measuring the NanoLuc (Nluc) signal in the feces of each mouse (E). Throughout the experiment, Nluc expression was measured daily in the feces of two groups of 5 GKO mice infected for 3 days with either *Cp*-INRAE-Nluc-mCh or *Cp*-INRAE-Nluc-CPSF3^Y385N^. The mice received 30 mg/kg AN3661 supplemented in their drinking water from 3 days post-infection onward. Starting from day 11 post-infection, both groups were switched to receive AN7973 at 15 mg/kg in their drinking water instead of AN3661. SEM images of the ileum showing total clearance in both groups treated with AN7973 (F).

We generated this transgenic *C. parvum* line by replacing the endogenous *CpCPSF3* locus with a *Cryptosporidium* codon-optimized transgene encoding *CpCPSF3* carrying the Y385N substitution. The repair cassette was designed to include the actin 3′ untranslated region (UTR) to ensure proper transcriptional termination of *CpCPSF3*, as well as an expression cassette encoding a nanoluciferase (Nluc)-neomycin phosphotransferase (Neo) fusion, separated by a self-cleaving P2A peptide, and placed under the control of the constitutively active enolase promoter (**Fig2-B**).

To generate the transgenic strain, the homologous repair (HR) template and a plasmid expressing Cas9 and the guide RNA (gRNA) were cotransfected. Following *in vivo* selection in GKO mice under paromomycin selection, the strain was purified, and correct insertion of the repair cassette was validated (**Fig2-C**). EC₅₀ values were then determined in infected HCT-8 for the three benzoxaboroles, using Halocur™, the reference drug for ruminant cryptosporidiosis that targets prolyl-tRNA synthetase as a control. A marked increase in resistance was observed only for AN3661, with *Cp*CPSF3_Y385N_ strain displaying an average 82-fold increase in resistance compared to the wild-type CPSF3-WT strain (**Fig2-D**). In contrast, no significant change in susceptibility was detected for AN7973 or AN13762 (**Fig2-D**). These results suggest that the Y385 residue is specifically critical for AN3661 binding to *Cp*CPSF3, whereas AN7973 and AN13762 either interact with *Cp*CPSF3 through distinct molecular contacts or act via an alternative target.

To confirm *in vivo* resistance to AN3661, we established a mouse model in which the compound was administered via drinking water to susceptible GKO mice from three days post-infection onward. We first determined that a dose of 20 mg/kg/day was sufficient to effectively control the *Cp*-INRAE-Nluc-mCh strain infection in GKO mice (**Fig2-E**). To ensure robust drug exposure, we increased the dose to 30 mg/kg/day and treated two groups of GKO mice starting from 3 days post-infection onward, one infected with the *Cp*-INRAE-Nluc-mCh strain and the other one with the *Cp*CPSF3_Y385N_ strain. A rapid decrease in Nluc activity was observed in the *Cp*CPSF3 strain, whereas activity continued to increase in the *Cp*CPSF3_Y385N_ mutant. Replacing AN3661 with AN7973 in the drinking water resulted in rapid control of the *Cp*CPSF3_Y385N_ strain and sustained control of the *Cp*CPSF3 strain (**Fig2-F**), confirming distinct mechanisms of action for the two compounds.

### AN7973 inhibits *T. gondii* growth independently of *Tg*CPSF3 amino acid substitutions, including Y483N, that confers resistance to AN3661 and AN13762

We next investigated whether AN7973 can inhibit *T. gondii* growth, similar to AN3661 and AN13762. Using the *T. gondii* RH *ku80* NLuc strain [10] (**Fig3**), we determined an EC₅₀ of ∼2 µM and CC_50_ of 193 µM, resulting in a SI of 97, in human foreskin fibroblasts (HFF). These results indicate that AN7973 is also a promising candidate for the treatment of toxoplasmosis. Since we previously generated *Tg*CPSF3 mutants resistant to AN3661 [7] and AN13762 [10], we used these strains to assess whether six key *Tg*CPSF3 substitutions (Y328C, Y483N, E545K, Y328H, G456S, and S519C) confer cross-resistance to AN7973. None of the mutants exhibited significantly elevated EC₅₀ values compared to the wild-type strain, strongly suggesting that AN7973 either acts differently on *Tg*CPSF3 or targets an additional factor (**Fig3**).

**Figure 3:**
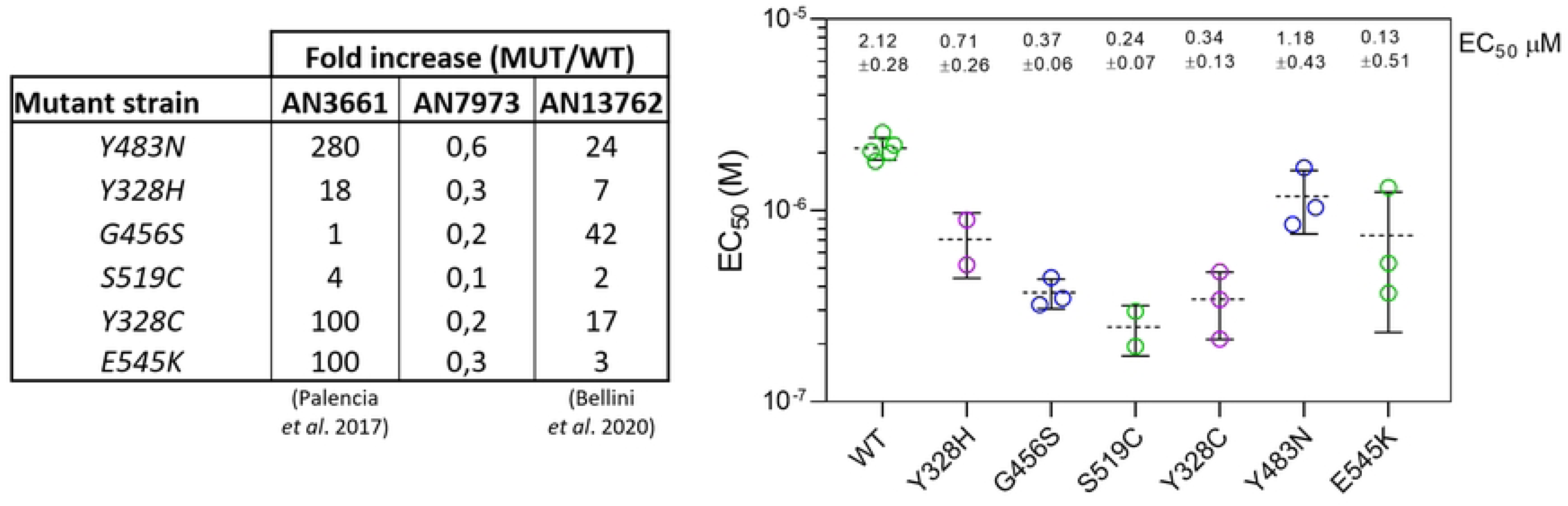
AN3661 and AN13962 *T. gondii* mutant strains resistance to AN7973. Resistance to AN7973 was assessed by comparing the wild-type RH *ku80* NLuc-P2A-EmGFP with six *T. gondii* clones known to be resistant to AN3661 and/or AN13762 (RH ku80 NLuc TgCPSF3^mut^). EC_50_ were determined on HFF infected cells exposed to various AN7973 concentrations. The Table show the fold increased EC_50_ values (EC_50mut_/EC_50WT_) for AN7973 and previously identified values for AN3661 and AN13762 [7, 10, 31]. EC_50_ individual values for AN7973 are shown against all six *T. gondii* mutant strains (right panel).

### Exposure of *C. parvum* to AN3661, AN7973 and AN13762 induce 3’ and 5’ overrunning transcripts

To dissect the mechanism of inhibition of the three benzoxaboroles on *Cryptosporidium*, we examined their impact on parasite pre-mRNA maturation. RNA was extracted from infected HCT-8 cells treated individually with the benzoxaboroles or left untreated as a control. Polyadenylated transcripts were enriched using polyT magnetic beads, reverse transcribed into mRNA/cDNA duplexes, and subjected to nanopore sequencing. Despite the low abundance of parasite RNA relative to host-cell RNA (1 to 5 %), a marked defect in mRNA end processing—affecting both 5’ ends and, more prominently, 3′ ends—was observed following treatment with AN3661, AN7973 and AN13762 (**Fig4-A**). At the gene level, it is evident that not all gene transcripts were directly affected, and many of the apparent 5′ defects likely correspond to 3′ readthrough events extending into downstream genes (examples shown in **Fig. 4B**). Among all tested conditions, AN3661 induces the most pronounced 3′ readthroughs, despite yielding the lowest number of parasite mRNA reads. This likely reflects skipping of canonical 3′ cleavage sites, triggering nonsense-mediated decay, with only transcripts polyadenylated at alternative downstream sites—often within adjacent sense genes—being captured. Interestingly, recently described polycistronic gene clusters in *Cryptosporidium* may be particularly susceptible to this phenomenon [19]. Notably, low levels of 3′ readthroughs are also detected in the untreated condition, likely reflecting the same underlying mechanism.

**Figure 4:**
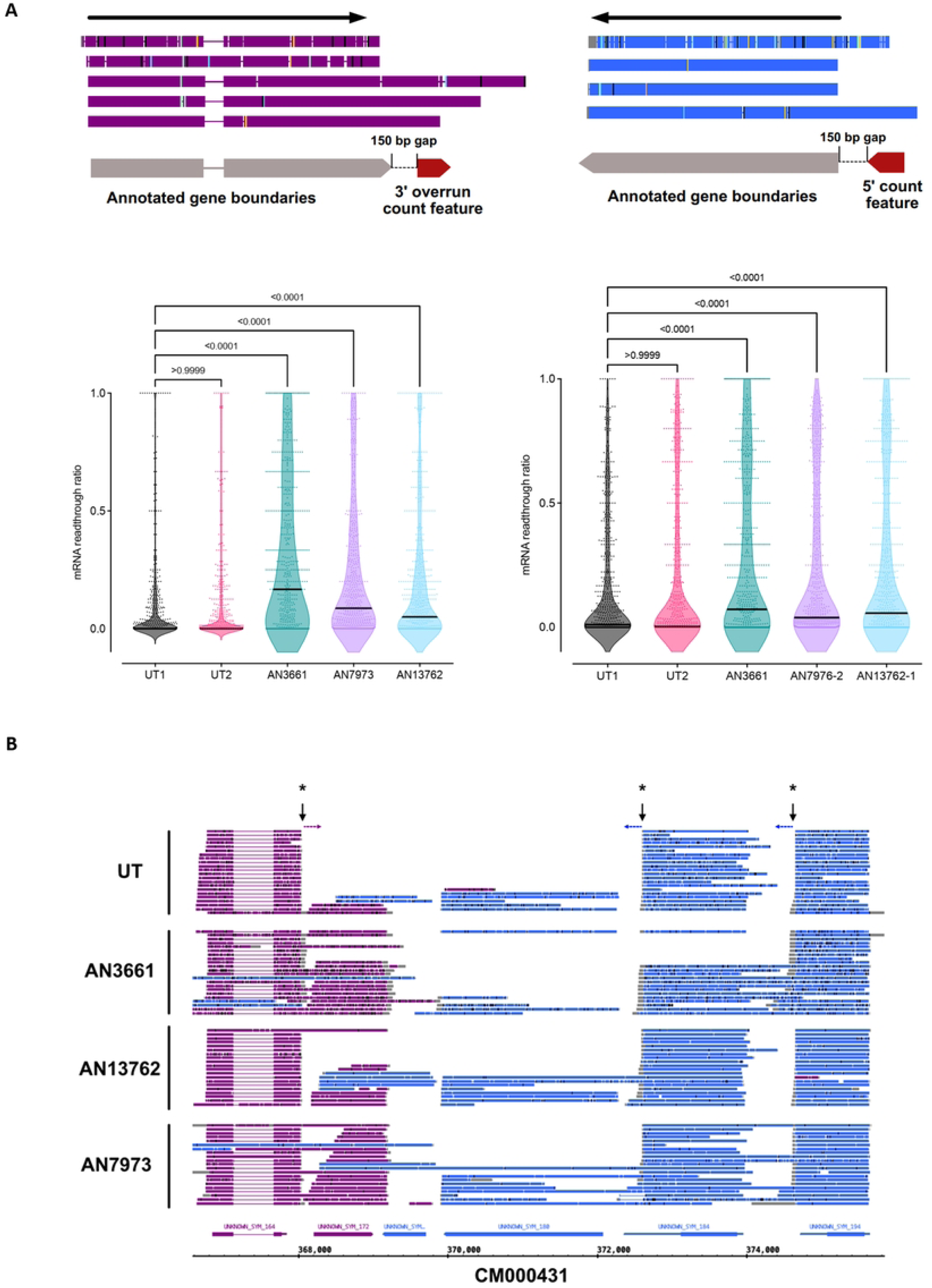
3′ and 5′ end quantification of readthroughs on mock or benzoxaborole-treated *C. parvum HCT-8 culture.* HCT-8 cells grown on a T25 flask were heavily infected with *C. parvum* oocysts (MOI=10) for 11h. After 1h oocysts were washed and cells treated with AN3661, AN7973, AN13762 with a concentration equivalent to 4-fold of their respective EC_50_ or mock treated. RNA was extracted and analyzed by nanopore. For genes with at least three detectable transcripts, readthrough sections 150 bp upstream (5′) and downstream (3′) were used as count features and divided by the total gene count to calculate a readthrough ratio (A). Statistics: Kruskal–Wallis multiple comparison tests were used to calculate the p-values indicated on the violon plot graphs below the graphic representations (A). Integrated Genome Browser snapshot within the CM000431 contig of multiple instances of readthroughs at the single molecule level. Readthrough sites are indicated by an arrow and asterix while the readthrough direction is shown with a dotted arrow (B).

Overall, these results demonstrate that AN3661, AN7973, and AN13762 all interfere with *Cryptosporidium* mRNA maturation, most likely by inhibiting the cleavage activity of *Cp*CPSF3.

### AN3661 inhibits *Eimeria tenella* and *Giardia duodenalis* development, while AN7973 inhibits *E. tenella*

Finally, we evaluated whether the two most efficient compounds against *Cryptosporidium*, AN3661 and AN7973, also exhibit activity against other protozoans of veterinary and public health importance. The critical Y385 amino acid for *Cryptosporidium* inhibition by AN3661 is also conserved in *Eimeria tenella* and *Giardia duodenalis* (**Fig5-A**). Using the *E. tenella* transgenic strain (*Et*-INRAE-Nluc-mCh) and the CLEC-213 chicken epithelial cell line infection model, we determined an EC_50_ of 43nM (SI > 325) with AN3661 and 86nM (SI > 140) with AN7973 (**Fig5-B**). Regarding *G. duodenalis*, variability in drug sensitivity between assemblages and isolates has already been documented [20, 21]. To account for these three isolates representing different genetic groups (assemblages) causing giardiasis in humans and/or livestock (WBC6, GS/M, P15) were tested. Trophozoite survival assays showed that all isolates were susceptible to AN3661, with EC₅₀ values ranging from20-40 µM (**Fig5-C**). These values were slightly higher compared to those of the reference drug the nitroimidazole metronidazole (MTZ, EC_50_ 5-10 µM, data not shown). For both parasites, CPSF3 amino acid sequences were found to be highly conserved among strains (**Fig5-D**). Together, these results extend the potential interest of AN3661 and AN7973 as candidates for a broader parasitic control.

**Figure 5:**
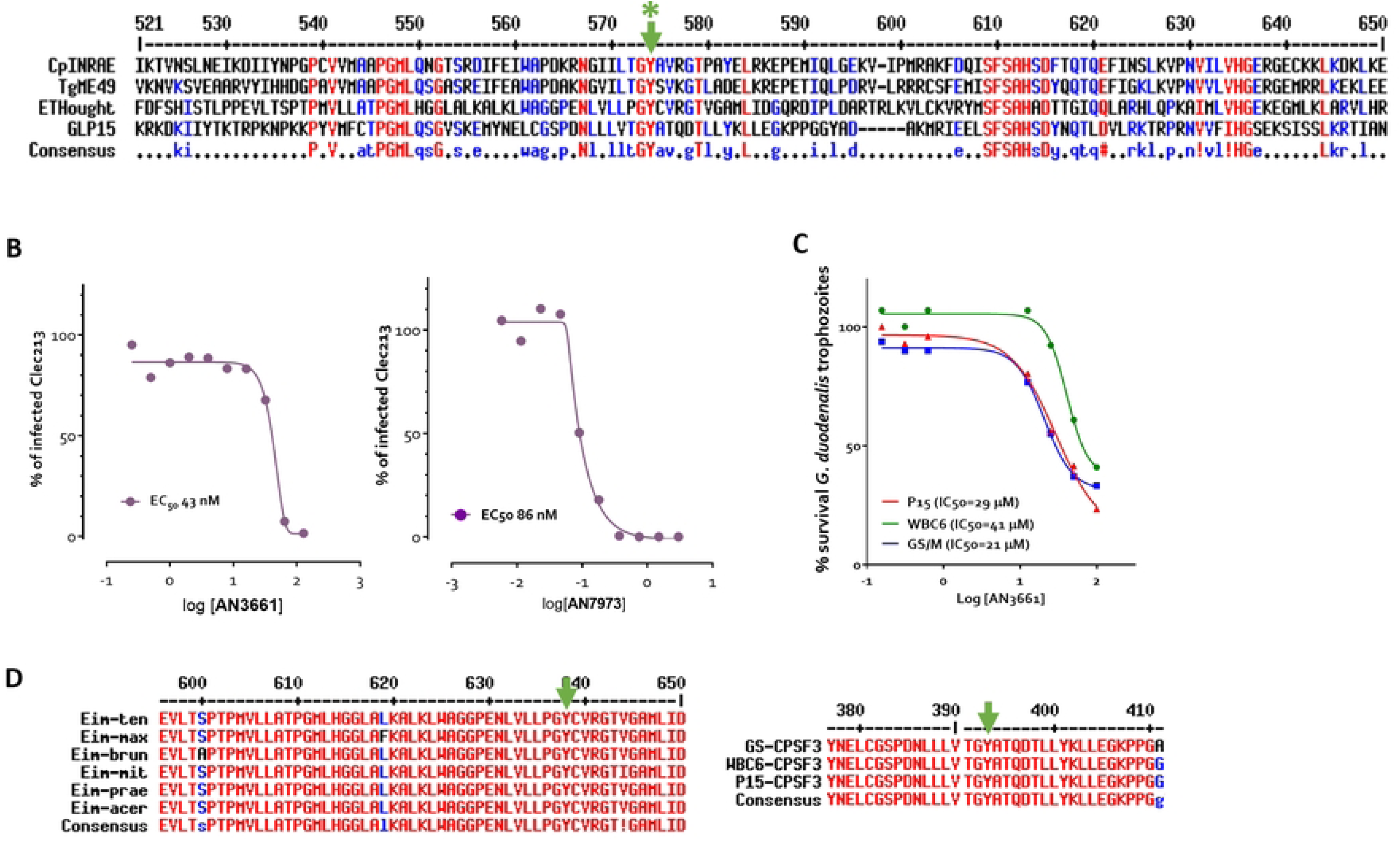
*In vitro* effects of AN3661 and AN7973 on *Eimeria* and *Giardia* development. Partial Amino acid alignment of CPSF3 from *C. parvum* INRAE strain, *T. gondii* ME49 strain, *E. tenella* Houghton strain and *G. lamblia/G. duodenalis* P15 strain (A). Alignment was performed with Multalin free software [32]. Green star and arrow show the conserved tyrosine (Y) amino acid that confers resistance to AN3661 in *T. gondii* and *C. parvum* (A). EC_50_ curves obtained by infecting CLEC-213 cells with *E. tenella* sporozoites expressing Nluc at a multiplicity of infection of 1 in the presence of various concentration of AN3661 or AN7973 (B). IC₅₀ determination for three *G. duodenalis* strains (P15, WBC6, GS/M) exposed to increasing concentrations of AN3661 (C). Amino-acid sequence alignment showing strong conservation in the region surrounding the tyrosine residue indicated by the green arrow among six chicken *Eimeria* species and three *Giardia* strains (D).

## Discussion

We previously demonstrated the efficacy of two orally administrable benzoxaboroles, AN3661 and AN13762, in controlling *C. parvum* infection in both neonatal and immunocompromised adult mouse models [6, 10]. Independently, Lunde *et al.* [14] reported the potent activity of a third benzoxaborole, AN7973, against *C. parvum*, although its proposed mechanism via inhibition of CPSF3 remained to be experimentally validated. While a crystallographic structure showed that AN3661 binds to *C. hominis* CPSF3, no reverse genetic evidence had yet identified resistance-conferring mutations in this gene in *Cryptosporidium*, and the exact mechanism of action, particularly the effect on 3’ end mRNA processing had not been fully investigated. In the present study, we leveraged orthologous mutations discovered in *T. gondii* and *P. falciparum* to confirm the role of CPSF3 in mediating the activity of AN3661, AN13762, and AN7973, three compounds suggested to share a common mode of action through CPSF3 inhibition.

Our results identify the 8-12h post-infection window as the most vulnerable stage for all three compounds, coinciding with the burst of mRNA synthesis required for first-generation merozoite formation. During this period, the compounds limit full development and egress of the merozoites *in vitro*. Incubation with host epithelial cells did not affect the infection process by sporozoites, and the drug withdrawal assays confirmed a parasiticidal effect on meronts with no parasite regrowth. These results are consistent with CPSF3 being a putative target of the compounds. Notably, the endonuclease CPSF3 is expressed all along the cell cycle of *Cryptosporidium* (data not shown). Using a qRT-PCR method with female and male markers we confirmed that AN3661 and AN7973 are also active to limit sex-stage development, whereas AN13762 does not. This lack of activity might reflect the inefficient conversion of AN13762 into its active metabolite AN10248, during sexual stages, as previously described in *Plasmodium* [8], thereby reducing its potency at this developmental stage *in vitro*. Indeed, although AN13762 controls *Cryptosporidium* effectively *in vivo*, it’s *in vitro* EC_50_ (∼10µM) [10]—over 100-fold higher than AN3661 [6]—likely reflects limited maturation into the fully active form under culture conditions.

All three drugs share a benzoxaborole core (Supplemental Fig1G) with AN7973 and AN13762 carrying longer side chains. To date, all the crystallized benzoxaboroles binding to CPSF3 rely primarily on the same key octahedral coordination between the boron atom and the two divalent cations in the active site (Supplemental Fig1A-1D) [6, 22]. These were originally believed to be zinc, however recent work on the human ortholog of CPSF3 (also known as CPSF73) has shown that these cations could also be a mixture of iron, manganese, zinc and nickel [23]. In any case, we can anticipate that all benzoxaboroles active against CPSF3 will likely bind similarly at least within the benzoxaborole moiety due to chemical coordination between the boron and zinc atoms enforcing a set of geometry constraints.

The crystal structure of *C. hominis* CPSF3 with AN3661 originally showed that the positioning of Y385 was adjacent to the AN3661 moiety (Supplemental Fig1A-B) and that its mutation into a asparagine likely either perturbed the hydrophobic binding to the compound without directly creating a steric clash, or had more of a allosteric effect by potentially affecting the mRNA substrate binding [10]. Since this conserved tyrosine residue was shown to be the most effective resistance conferring mutation in *Toxoplasma* and *Plasmodium* for AN3661 and AN13762 [7, 10, 15], we decided to generate a new *C. parvum* strain presenting a single substitution of the tyrosine to an asparagine (Y385N). Analogous to *T. gondii* and *P. falciparum*, this mutation also triggered an increased resistance to AN3661 of about 80-fold. Surprisingly though, the mutant strain exhibited similar sensitivity to AN13762, suggesting that the effect of this mutation is much stronger on a smaller molecule such as AN3661. Similarly, Y385N mutation does not increase resistance to AN7976. To determine whether all three compounds affect mRNA transcripts, we performed nanopore direct RNA sequencing on *C. parvum*-infected cells treated with benzoxaboroles or mock-treated controls. All three compounds affected *Cryptosporidium* transcripts with significant 3’overrun but also defects on 5’ with AN3661 which will result in alteration of protein synthesis. This process is performed by the CPSF complex machinery [24] therefore reinforcing the idea that AN7973 and AN13762 also bind and inhibit CPSF3. Using Alphafold 3 [25], we predicted that AN7973 and AN13762 can bind within the RNA binding groove only if the Apicomplexan specific insert loop exits the conformation observed within the AN3661 crystal structure (Supplemental Fig1B, 1E-F). As such, this extended loop in Apicomplexa which varies in between species may be one of the contributing factors of compound selectivity. Due to the extension beyond the benzoxaborole moiety, many other interactions come into play, which may explain why in *C. parvum*, the Y385N mutation has no effect on sensitivity in comparison to AN3661, which almost exclusively relies on the octahedral coordination on the divalent cations. In *Trypanosoma brucei*, AN7973 causes trans-splicing to fail within just an hour and the compound effectiveness is reduced when CPSF3 is over-expressed suggesting that CPSF3 is likely the molecular target [9]. In *Toxoplasma*, we showed for the first time that AN7973 limits parasite growth *in vitro*. However, none of the six mutants conferring resistance to AN3661 and AN13762 exhibited increased resistance to AN7973 compared to the parental strain, suggesting a distinct binding site, or alternatively, an interaction with a different target involved directly or indirectly in mRNA processing. This indicates that, despite the high conservation of CPSF3 among apicomplexans, variations in the binding site may exist, and functional validation is essential before extrapolating results across the group. AN3661 and AN13762 are alternative chemotypes in their binding to *Tg*CPSF3. Here, we demonstrated that AN7973 can rapidly eradicate the AN3661-resistant strain (*Cp*CPSF3_Y385N_) in heavily infected GKO mice. This is of high importance to face parasite resistance by offering the possibility to combine drugs or use them in rotation to limit occurrence of resistance.

In *Cryptosporidium* resistance to current low-efficacy treatments has not been reported, yet. However, Hasan *et al.* documented spontaneous, clinically relevant drug resistance to the methionyl-tRNA synthetase inhibitor 2093 in calves [26]. Although this inhibitor was initially highly effective in controlling *Cryptosporidium* development in young calves within a few days, the emergence of a resistant strain was observed in several animals. This strain carried point mutations in the *MetRS* gene, and further investigation confirmed that these mutations were responsible for the resistance. This highlights the need to anticipate potential anticryptosporidial resistance and to develop strategies such as combination of alternative chemotypes on defined targets.

Furthermore, AN3661 was found to be effective in inhibiting *G. duodenalis* trophozoite proliferation as well as the intracellular development of *E. tenella*, the latter also being susceptible to AN7973. Coccidiosis, caused by *Eimeria spp*., leads to substantial financial losses in poultry production [27]. Giardiasis affects approximately 280 million people annually, making it one of the most prevalent parasitic gastrointestinal diseases worldwide [28]. Giardiasis in livestock can cause mild to moderate disease, mainly associated with diarrhea, particularly in young animals. Although the economic burden of *G. duodenalis* infection in livestock is poorly described, the easy spread of the disease among farm animals and the growth impairment can lead to lower carcass weight and dressing percentages at slaughter thus potentially cause considerable economic losses on farms worldwide [29]. Therefore, identifying new drug treatments is of great importance for controlling these diseases both in human and animals. However, it remains to be determined if CPSF3 is the target of these compounds in *Giardia* and *Eimeria spp.*.

CPSF3 is a highly conserved, essential endonuclease in apicomplexan parasites, making it a particularly attractive drug target. In *Cryptosporidium*, CPSF3 has shown strong potential for controlling cryptosporidiosis. Its high level of conservation across species further supports its relevance for the development of broad-spectrum antiparasitic therapies. Its expression throughout the *Cryptosporidium* cell cycle and its involvement in critical processes such as mRNA maturation make this endonuclease a target of choice. Orally administrable compounds like AN3661, AN7973, and AN13762 efficiently inhibit the parasite by interfering with mRNA processing, though their binding selectivity may vary among species. Combining such compounds represents a promising strategy to limit the emergence of resistance. Here, we provide functional evidence that AN3661 targets CPSF3, with the amino acid residue Y385 playing a key role to counter its effect. In contrast, AN7973 and AN13762 sensitivity in *Cryptosporidium* seems unaffected by this commonly found mutation in other species. Finally, we demonstrate that these compounds are active against two other apicomplexan parasites of veterinary and public health importance, namely *Eimeria* and *Giardia*, respectively.

## Materials and Methods

### Parasites

The wild-type and transgenic strains of *Cryptosporidium*, *Toxoplasma*, *Eimeria*, and *Giardia* used in this study are listed in Supplementary Table 1. *Cryptosporidium* strains were maintained by serial passage in GKO mice infected with 5 × 10⁵ oocysts. *Toxoplasma gondii* was propagated *in vitro* through serial passage on monolayers of human foreskin fibroblasts (HFF). *Eimeria tenella* was maintained by passage in SOPF outbred PA12 Leghorn chickens infected with 10⁴ sporulated oocysts. Axenic cultures of *G. duodenalis* trophozoites of isolates WB clone C6 (WBC6, ATCC 50803, Assemblage AI) and GS/M (ATCC 50581, Assemblage BIV) and P15 (Assemblage E, kindly provided by Prof. Staffan Svärd, Uppsala University, Sweden) were grown in screw-cap tubes (NuncTM, ThermoFisher Scientific, Waltham, MA USA) in 10 ml of TYI-S33 medium supplemented with 10% adult bovine serum (Euroclone, Italy); 100 mg/mL streptomycin and 100 U/mL penicillin (Euroclone, Italy) and bovine bile (Sigma-Aldrich S.r.l., Italy) (Keister 1983) at 37°C until confluence for 48-72 hours (h). Cryptosporidium parasites were purified as previously described. [30]. For *Eimeria tenella*, unsporulated oocysts were harvested from the caeca and purified using filtration, magnesium sulfate flotation, and a discontinuous sucrose gradient. Sporulation was achieved by incubating oocysts in a 2.5% potassium dichromate solution (Sigma-Aldrich) at 27°C for 72 hours under agitation in a water bath. Sporozoites were then purified from sporulated oocysts as follows: oocyst walls were disrupted using 0.5 mm glass beads, and the released sporocysts were incubated at 41°C for 1 hour in a solution containing 0.25% porcine trypsin (Sigma-Aldrich) and 0.5% porcine bile salts (Sigma-Aldrich) in PBS (pH 7.4). Finally, sporozoites intended for in vitro infection experiments were purified by sequential filtration through cotton and 5 µm polycarbonate filters (Whatman).

### Cell culture

Human ileocecal adenocarcinoma cells (HCT-8 cells; ATCC CCL-34) were cultured in RPMI 1640 medium supplemented with glutamine (Gibco), 10 % fetal bovine serum (Dutscher), 50U/ml penicillin, 50 µg/ml streptomycin (Gibco), 15 mM HEPES (Gibco, HEPES Buffer Solution 1 M), and 1 x Sodium pyruvate (Gibco, Sodium pyruvate 100 mM, 100). Human primary fibroblasts (HFFs, ATCC® CCL-171™) were cultured in Dulbecco’s Modified Eagle Medium (DMEM) (Invitrogen) supplemented with 10% heat inactivated Fetal Bovine Serum (FBS) (Invitrogen), 10 mM (4-(2-hydroxyethyl)-1-piperazine ethanesulphonic acid) (HEPES) buffer pH 7.2, 2 mM L-glutamine and 50 μg/ml of penicillin and streptomycin (Invitrogen). CLEC213 chicken epithelial cells were cultured in Dulbecco’s modified Eagle’s medium F12 (DMEM F12) medium supplemented with 10% fetal bovine serum (FBS, Dutscher), penicillin (50 U/ml, Cytiva, Hyclone), and streptomycin (50 µg/ml, Cytiva, Hyclone). The cultures were free of mycoplasma, as determined by qualitative PCR and cells were incubated at 37°C in 5% CO_2_ except for CLE213 incubated at 41°C in 5% CO_2_.

### *In vitro* infection and EC_50_ determination

#### Cryptosporidium

Human ileocecal adenocarcinoma cells (HCT-8) cells were grown to confluence in white 96-well plates (Thermo Scientific Nunc MicroWell), infected with freshly purified *Cp*-INRAE-Nluc-mCh-expressing oocysts at a multiplicity of infection of 0.5 for three hours. After three washes in PBS the compounds were added at different concentrations (6 replicates for each concentration) and cells were incubated for 48h at 37°C. Three third of the culture supernatant was removed from the wells and 25 µl of Nano-Glo® Luciferase assay buffer containing 1:50 of Nano-Glo® Luciferase assay substrate (Promega) was added to the wells. After 3 min of incubation, luminescence was measured with the GloMax® Explorer multimode microplate reader (Promega). EC_50_ were calculated with GraphPad Prism 6 software from the asymmetric sigmoidal, 5PL curve.

#### Toxoplasma

HFF cells were grown to confluence in 96-well plates and infected for 2h with 2,000 tachyzoites of *T. gondii* RH ku80 NLuc-P2A-EmGFP strain or TgCPSF3 mutant strains (RH *ku80 NLuc TgCPSF3^mut^*), all expressing the nanoluciferase. AN7973 was diluted in growth medium and added to the monolayers at various concentrations in triplicates along with DMSO-treated controls. At 48hpi, the NanoLuc assays were performed using the Nano-Glo® Luciferase Assay System according to manufacturer’s instructions (Promega) and previously published protocol [10]. EC_50_ were determined using non-linear regression analysis of normalized data and assuming a sigmoidal dose response. EC_50_ values for each compound represent an average of at least two independent biological replicates. AN7973 cytotoxicity was assayed on HFF cells after 72 h of incubation using CellTox^TM^ Green Cytotoxicity Assay (Promega).

#### Eimeria

CLEC-213 cells were seeded in 96-well plates at 1.5x10^4^ cells per well and infected the next day with Et-INRAE-Nluc-mCh freshly excysted sporozoites preincubated with the compounds for 1h. Seventy-two hours later, parasite development was assessed using the Nano-Glo® luciferase assay (Promega). Values are sextuplicate for each drug concentration ± SEM. EC_50_ were calculated with GraphPad Prism 6 software from the asymmetric sigmoidal, 5PL curve. The half-maximal cytotoxic concentration (CC50) was determined by adding the compound on top of CLEC-213 cells monolayer at different concentrations for 72h at 41°C. To quantify cytotoxicity, MTS was added following manufacturer’s instructions (CellTiter 96® AQueous One Solution Cell Proliferation Assay, Promega) and cells were incubated at 37°C for 4h. Absorbance was read at 490 nm on a microplate reader (GloMax® Explorer, Promega).

#### Giardia

Axenic cultures of *G. duodenalis* grown at confluence for 48-72 hours in 96-well microplates format were used to evaluate drug susceptibility as previously described [21]. Trophozoites from a log-phase culture were harvested by chilling on ice and counted in hemocytometer (KovaTM, Thermo Fischer Scientific, Waltham, MA, USA). Trophozoites were seeded in 200 μl of modified TYI-S-33 medium at 0.5x10^5^ parasites/well for WBC6 and GS/M and at 0.8x10^5^ parasites/well for P15. AN3661 (HY-128204) and metronidazole (HY-B0318) 10 mM stock solutions in DMSO were purchased from MedChemExpress (D.B.A. ITALIA S.R.L., Milan, Italy). Two-fold serial dilutions of the compounds were prepared in DMSO at 100X concentration and individually added to each well. Parasites were incubated at 37°C for 48 hours in anaerobic conditions by placing microplates in Oxoid™ AnaeroGen™ Compact Sachet (Thermo Scientific™). After 48h, trophozoites viability was determined by the bioluminescent ATP content assay, according to the manufacturer’s instructions (CellTiterGlo 2.0, Promega Italia, Milan, IT). Luminescence was read on a microplate reader (GloMax® Explorer, Promega). were calculated with GraphPad Prism 6 software from the asymmetric sigmoidal curve. Each experiment was done in triplicate and at least three biological replicates were performed.

### Q-RT-PCR for evaluating sexual development

DMC1 and AP2-M genes have been selected because of their selective expression for female and male gametes. The following primers were used for Q-RT-PCR analyses: CpDMC1_Fw 5’GTTGATGGGCGGATTTGAAAG3’; CpDMC1_Rv 5’AACAGACTTTCCCATTACCTTCC3’; AP2-M_Fw 5’ATACGTCGGATGGGTTGCTT3’; AP2-M_Fw 5’GCTCCCATGCCTCTATTTCC3’. RNA from infected cells were collected in Trizol^TM^ and extracted and purified with a kit Zimo Direct-Zol^TM^ RNA microprep including DNAse treatment. A second DNAse treatment was performed with RNA Clean and Concentrator-5^TM^ before reverse transcription with M-MLV Reverse transcriptase (Promega) in presence of RNasin® Ribonuclease Inhibitor. 2 ^−ΔΔCt^ relative quantity was determined using Human GAPDH as the reference housekeeping gene.

### Ethics statement

Animal needs were met in accordance with the European Community Council Directive 2010/63/EU. The experimental facilities had received authorization to house experimental animals from the local bureau of veterinary services (Indre-et-Loire, France, authorization no. D 37-175-3), and all the experimental procedures were approved by the Val de Loire Ethics Committee (CEEA19) (authorization no. APAFlS#50777 and #27679). All the personnel involved had dedicated training in animal care, handling, and experimentation, as required by the French Ministry of Agriculture.

### *In vivo* experiment and parasite load determination

All animal experimentations have been performed in the “Infectiology of Farm, Model and Wildlife Animals Facility” (PFIE, Centre INRAE Val de Loire: doi.org/10.15454/1.5572352821559333E12; a member of the National Infrastructure EMERG’IN: doi.org/10.15454/90CK-Y371). Specific-pathogen-free (SPF) C57BL/6J interferon-gamma-deficient (IFN-γ^-/-^: GKO) mice were individually marked and housed in groups in cages on poplar sawdust bedding. AN3661 or AN7973 was was administered via the drinking water. Mice were orally infected with 5 × 10⁶ oocysts of the *Cryptosporidium parvum Cp*-INRAE-Nluc-mCh strain, which allows quantification of parasite load throughout the experiment. For individual fecal sample collection, mice were briefly placed in separate boxes to ensure accurate sampling. For Nluc signal determination, fecal samples were collected and weighed. Each sample was placed in a tube containing ten 3mm glass beads and 500 µL of fecal lysis buffer composed of: 50 mM Tris-HCl (pH 7.5; stock 0.5 M), 2 mM dithiothreitol (DTT), 2 mM EDTA, 10% glycerol, and 1% Triton X-100. Samples were vortexed for 30 seconds, then incubated for 30 minutes at 4 °C. Following incubation, samples were vortexed again for 30 seconds and centrifuged for 1 minute at 10,000 rpm. From each sample, 50 µL of supernatant was transferred to a white 96-well plate. An equal volume (50 µL) of Nano-Glo lysis buffer containing Nano-Glo substrate (Promega, Nano-Glo Luciferase Assay System, ref. N1130) diluted 1:50 was added to each well. After a 3-minute incubation at room temperature, luminescence was measured using a GloMax-Multi Detection System.

### Timelapse and electron microscopy

Timelapse imaging was performed using a Nikon AX R confocal microscope equipped with a 60×/1.2 NA water-immersion objective, a temperature-controlled incubation chamber (5% CO₂, 37 °C), and a programmable water dispenser. Multidimensional acquisition was automated using the JOBS module of the NIS-Elements AR software (version 5.42.07, Nikon). Samples were cultured in 8-well ibidi chambers and imaged every 15 minutes. For each condition, three positions were selected, and z-stacks were acquired. Images were subsequently extracted, compiled using FIJI (version 1.5), and manually analyzed. Scanning electron microscopy (SEM) was performed on ileal tissue samples from euthanized, *Cryptosporidium*-infected mice treated sequentially with AN3661 and AN7973. Samples were fixed for 24 hours in 4% paraformaldehyde and 1% glutaraldehyde in 0.1 M phosphate buffer (pH 7.3), and processed as previously described [10]. SEM imaging was conducted using a Zeiss Ultra Plus field emission gun scanning electron microscope (FEG-SEM).

### Generating point mutation in *Cryptosporidium*-CPSF3 using CRISPR/cas9: To generate the *C. parvum*

CPSF3_Y385N_ mutant strain, excysted oocysts were electroporated with the plasmid pACT1-Cas9-GFP-U6::CpCPSF3 (GenBank Submission # PX246137) together with a repair DNA fragment carrying the *Cp*CPSF3^Y385N^ mutation. The pACT1-Cas9-GFP-U6::CpCPSF3 plasmid encodes a single guide RNA (sgRNA) targeting a 20 bp sequence located 12 bp upstream of the desired editing site. The DNA repair fragment was amplified by PCR using primers HR-CpCPSF3_F (5′-GAAATTAAAGATATTATCTATAATCCAG-3′) and HR-CpCPSF3_R (5′-CAAGTCATGAAAAGGCTAGAGAAGC-3′), with pCPSF3^Y385N^-Nluc-P2A-NeoR-CPSF3 (GenBank Submission # PX246138) as template. Both the pACT1-Cas9-GFP-U6::CpCPSF3 and the pCPSF3^Y385N^-Nluc-P2A-NeoR-CPSF3 vectors were DNA-synthesized by GenScript. Note that the PAM sequence was mutated in the repair template to prevent Cas9-mediated further cleavage.

## Acknowledgements

We are very grateful to the PFIE technical staff for their assistance with animal experimentation and for ensuring compliance with ethical and experimental standards. We also thank P-I Raynal from the Microscopy Division of Tours-university for SEM pictures. This work was supported by the Agence Nationale pour la Recherche (ANR) fundings ApiNewDrug (ANR-21-CE35-0010-0 to F.L. and M.A.H), and ANR-11-LABX-0024 via the Laboratoire d’Excellence (LabEx) ParaFrap. The Labex also funded PhD project of J. W.. This work was also supported by the European Partnership on Animal Health and Welfare (PAHW), co-funded by the European Union under Horizon Europe, Grant Agreement No. Grant Agreement No 101136346 (SOA19 action to F.L. and M. L.).

## Author Contributions

F.L, A.B. and M.A.H. conceived the project and supervised the research. F.L., A.B., C.S., L.B., M.A.H., L.B., J.W., I.D, J.P. designed and interpreted the experimental work. J.W., L.B., C.S., A.G., J.P., L.S., T.B., I.D., M.L. performed the experiments. FL wrote the paper with editorial support from C.S., M.A.H., A.B., L.B and M.L.

## Supporting information captions

**Supplementary Figure 1:** Benzoxaborole binding against *Cryptosporidium* and Human CPSF3. (A) Overall view of *Ch*CPSF3 shown in orange bound to AN3661 shown in turquoise. (B) Zoomed in view of the *Ch*CPSF3/AN3661 structure with the Y385 residue highlighted in magenta and the Apicomplexan insert loop indicated which bares the E135 residue. (C) Structurally aligned view of *Hs*CPSF3 bound to the benzoxaborole cmp1. (D) Structurally aligned view of HsCPSF3 bound to the benzoxaborole cmp2. (E) Structurally aligned view of ChCPSF3 bound to AN7976 AlphaFold3 modelling. (F) Structurally aligned view of ChCPSF3 bound to AN13762 AlphaFold3 modelling. (G) 2d chemical diagram of the benzoxaboroles used in this study.

**Supplementary Table 1**: Parasite strains used in this study. Light blue domain is the conserved boron-containing heterocycle between the three compounds.

## Notes

### Competing Interest Statement

The authors have declared no competing interest.

